# The Draft Genome of *Kochia scoparia* and the Mechanism of Glyphosate Resistance via Transposon-Mediated *EPSPS* Tandem Gene Duplication

**DOI:** 10.1101/600072

**Authors:** Eric L. Patterson, Christopher A. Saski, Daniel B. Sloan, Patrick J. Tranel, Philip Westra, Todd A. Gaines

**Author notes:** Corresponding author: Todd Gaines, Department of Bioagricultural Sciences and Pest Management, Colorado State University, Fort Collins, CO 80523, USA. **Data deposition** All raw sequence read files for the whole genome sequencing have been deposited in the Sequence Read Archive database at NCBI under BioProject ID PRJNA526487 (SRR8835960-SRR8835963). The genome assembly was submitted to the NCBI genomes database with the accession SNQN00000000.

## Abstract

Increased copy number of the 5-enolpyruvylshikimate-3-phosphate synthase (*EPSPS*) gene confers resistance to glyphosate, the world’s most-used herbicide. There are typically three to eight *EPSPS* copies arranged in tandem in glyphosate-resistant populations of the weed kochia (*Kochia scoparia*). Here, we report a draft genome assembly from a glyphosate-susceptible kochia individual. Additionally, we assembled the *EPSPS* locus from a glyphosate-resistant kochia plant by sequencing a kochia bacterial artificial chromosome library. These resources helped reconstruct the history of duplication in the structurally complex *EPSPS* locus and uncover the genes that are co-duplicated with *EPSPS*, several of which have a corresponding change in transcription. The comparison between the susceptible and resistant assemblies revealed two dominant repeat types. We discovered a FHY3/FAR1-like mobile genetic element that is associated with the duplicated *EPSPS* gene copies in the resistant line. We present a hypothetical model based on unequal crossing over that implicates this mobile element as responsible for the origin of the *EPSPS* gene duplication event and the evolution of herbicide resistance in this system. These findings add to our understanding of stress resistance evolution and provide an example of rapid resistance evolution to high levels of environmental stress.

## INTRODUCTION

Gene copy number variation is an important source of genetic variation that can be deleterious in some cases, such as causing cancer in humans, and that can also increase genetic variation and lead to adaptations (Schimke et al. 1985; Lynch and Conery 2000; DeBolt 2010; Xi et al. 2011; Hull et al. 2017). This is especially true in plants where novel genetic variation is essential in the face of rapidly changing environments (DeBolt 2010). Increases in copy number of the 5-enolpyruvylshikimate-3-phosphate synthase (*EPSPS*) gene can confer resistance to glyphosate, the world’s most-used herbicide, in several plant species (reviewed in Sammons and Gaines 2014). These increases result in the over-production of the EPSPS protein, glyphosate’s target (Gaines et al. 2010; Wiersma et al. 2015), making it necessary for the application of more glyphosate to have the same lethal effect (Vila-Aiub et al. 2014; Godar et al. 2015; Gaines et al. 2016; Koo et al. 2018). This phenomenon has been observed in eight weed species to date; however, the DNA sequence surrounding the *EPSPS* gene duplication has only been resolved in one species, *Amaranthus palmeri* (Molin et al. 2017; Patterson et al. 2018), as most weed species do not have sequenced genomes. In the case of *A. palmeri*, *EPSPS* gene duplication is caused by a large, circular, extra-chromosomal DNA element that replicates autonomously from the nuclear genome (Molin et al. 2017; Koo et al. 2018). This mechanism results in *A. palmeri* plants containing up to hundreds of *EPSPS* copies (Gaines et al. 2010).

Recently, *EPSPS* gene duplication has been described in the weed species *Kochia scoparia* (kochia, syn. *Bassia scoparia*), one of the most important weeds in the Central Great Plains of the United States and Canada (Beckie et al. 2013; Jugulam et al. 2014; Beckie et al. 2015; Kumar et al. 2015; Wiersma et al. 2015; Gaines et al. 2016; Martin et al. 2017; Beckie et al. 2018). In glyphosate-resistant kochia, *EPSPS* copy numbers typically range from 3 to 8 with the highest reports at 11 copies (Gaines et al. 2016). In contrast to the extrachromosomal element observed in *A. palmeri*, fluorescence *in situ* hybridization (FISH) has shown that the *EPSPS* copies in kochia are arranged in tandem at a single chromosomal locus and are most likely generated by unequal crossing over (Jugulam et al. 2014). More detailed cytogenetics studies using fiber-FISH estimated that most repeats of the *EPSPS* loci are either 45 kb or 66 kb in length. Both inverted repeats and repeats of 70 kb in length were also observed (Jugulam et al. 2014). The initial event that started *EPSPS* gene duplication, the fine-scale sequence variation between the various types of repeats, and the other genes that may be co-duplicated with *EPSPS* remain unresolved.

Understanding how gene copy number variants form and their potential phenotypic consequences is essential for determining how plants adapt to their environment and thrive in adverse conditions. In this paper, we sequenced and assembled the genome of a glyphosate-susceptible kochia plant. We then identified the contig containing the *EPSPS* locus and investigated the genes that are co-duplicated with *EPSPS*, their transcription in glyphosate resistant and susceptible plants, and through whole-genome resequencing of a glyphosate-resistant plant, discovered the upstream and downstream borders of the duplicated region. We next sequenced and assembled the *EPSPS* locus from a glyphosate-resistant kochia plant using bacterial artificial chromosomes (BACs) probed for 1) the *EPSPS* gene, 2) the downstream junction, and 3) the upstream junction. After assembling four BACs we generated a model sequence of the *EPSPS* duplicated locus containing six instances of the *EPSPS* gene. We discovered two dominant repeat types with occasional inversions and repeats of different sizes using a combination of qPCR markers, genomic resequencing, and RNA-Seq data. Through this analysis, we also discovered a 16 kb mobile genetic element (MGE) that is associated with the gene duplication event. This MGE contains four putative coding sequences. We hypothesize that the insertion of this MGE downstream of the *EPSPS* gene is responsible for a disruption of this region and the origin of the *EPSPS* gene duplication event.

## MATERIAL AND METHODS

### Tissue Collection and Nucleic Acid Extraction

The herbicide-susceptible *K. scoparia* line “7710” (Preston et al. 2009; Pettinga et al. 2018) was used for genomic sequencing. All plants in this line were consistently controlled by glyphosate treatments at field rates of 860 g a.e. ha^−1^. Plants were grown in a greenhouse at Colorado State University. After seeds germinated, they were transferred into 4 L pots filled with Fafard 4P Mix supplemented with Osmocote fertilizer (Scotts Co. LLC), regularly watered, and grown under a 16-hr photoperiod. Temperatures in the greenhouse cycled between 25 ℃ day and 20 ℃ nights. A single, healthy individual was selected for tissue collection.

A glyphosate resistant line (M32) was obtained from a field population near Akron, Colorado (40.162382, −103.172849) in the autumn of 2012. After glyphosate failed to control these plants in the field, seed was collected from ten surviving individuals. Seeds were germinated and treated with 860 g a.e. ha^−1^ of glyphosate and ammonium sulfate (2% w/v). Survivors were then collected, crossed and seed was collected. This process was repeated for three generations until no susceptible individuals were observed in the progeny. All plants were confirmed to have elevated *EPSPS* copy number using genomic qPCR (Gaines et al. 2016).

For shotgun genome Illumina sequencing of the two lines, DNA was extracted from samples using a modified CTAB extraction protocol (see Supporting Information). For large-fragment, genomic PacBio sequencing of the glyphosate-susceptible line, the CTAB protocol was further modified to obtain more DNA of sufficiently large size (>10kb) (see Supporting Information). For RNA-Seq, susceptible and resistant plants were grown in the greenhouse as described above, until they were ~10 cm tall and 100 mg of young expanding leaf tissue was taken from each plant. RNA was extracted from young leaf tissue from four plants from each of the glyphosate-susceptible and resistant lines using the Qiagen RNeasy Plus Mini Kit. Each replicate sample was normalized to a total mass of 200 ng total RNA.

### Sequencing Libraries

Three genomic DNA libraries of glyphosate-susceptible kochia DNA were prepared for Illumina sequencing on a HiSeq 2500 at the University of Illinois, Roy J. Carver Biotechnology Center for genome assembly. First, DNA was size selected to 240 bp so that there was overlap between the read pairs in a high-coverage, short-insert library sequenced on one full flow cell (8 lanes) for use with ALLPATHS-LG. Second, two large insert, mate-pair libraries (5 kb and 10 kb) were each run on 1 lane at 2×150 bp.

Additionally, genomic DNA from the glyphosate resistant line was prepared for Illumina sequencing using the Genomic DNA Sample Prep Kit from Illumina following the manufacturer’s protocols and sequenced on one entire lane of a HiSeq 2500 flow cell. Quality of the raw Illumina sequence reads was assessed using FASTQC v0.10.1. Adapters were removed using Trimmomatic version 0.60 with the parameters “ILLUMINACLIP: tranel_adaptors.fa:2:30:10 TRAILING:30 LEADING:30 MINLEN:45” using these adapters: “AGATCGGAAGAGCAC” and “AGATCGGAAGAGCGT”.

A large insert DNA library for PacBio sequencing was generated at the UC Davis Genome Center using the PacBio SMRT Library Prep for RSII followed by BluePippin size selection for fragments >10 kb. The library was sequenced with 12 PacBio SMRT cells using the RSII chemistry after a titration cell to determine optimal loading. In total, 2,760,348 PacBio reads were generated with a read N50 of 6,576 bp with the largest read being 41,738 bp.

Strand-specific RNA-Seq libraries were prepared robotically on a Hamilton Star Microlab at the Clemson University Genomics and Computational Facility following in-house automation procedures generally based on the TruSeq Stranded mRNAseq preparation guide. The prepared libraries were pooled and 100 bp paired-end reads were generated using a NextSeq 500/550.

### Susceptible Genome Assembly

Two different assemblies were generated that integrated the PacBio and Illumina data of the susceptible kochia line. These two assemblies were then compared and merged by consensus for a single final assembly referred to as KoSco-1.0. For the first assembly, raw PacBio reads were error corrected using the high coverage, paired-end Illumina library with the error correcting software Proovread 2.13.11 (Hackl et al. 2014). Proovread was run with standard parameters, using the high coverage 150 bp, paired-end Illumina library on each SMRT cell individually. Error corrected reads were then assembled using the Celera Assembler fork for long reads, Canu 1.0 (Koren et al. 2017). Canu was run with a predicted genome size of 1 Gb, and the PacBio-corrected settings. For the second assembly, an initial ALLPATHS-LG v r52488 assembly was made with all three Illumina libraries (Butler et al. 2008). ALLPATHS was run assuming a haploid genome of 1 Gb. The resulting contigs were then scaffolded using the uncorrected PacBio reads and the software PBJelly 15.8.24 (English et al. 2012). PBJelly was run with the following blasr settings: -“minMatch 8 -sdpTupleSize 8 -minPctIdentity 75 -bestn 1-nCandidates 10 -maxScore −500 -nproc 19 –noSplitSubreads”. The two assemblies were then merged with GARM Meta assembler 0.7.3 to get a final version of the genome assembly for our analysis (Mayela Soto-Jimenez et al. 2014). The assembly from ALLPATHS was set to assembly “A” and the assembly from Canu was set as genome “B.” All other parameters were kept standard. We refer to the resulting meta-assembly as KoSco-1.0

### Genome Annotation

The merged assembly was annotated with the WQ-Maker 2.31.8 pipeline in conjunction with CyVerse (Cantarel et al. 2008; Thrasher et al. 2014). WQ-Maker was informed with kochia transcriptome from Wiersma et al. (2015), all expressed sequence tags (ESTs) from the Chenopodiaceae downloaded from NCBI, all protein sequence from the Chenopodiaceae family downloaded from NCBI, and Augustus using *Arabidopsis thaliana* gene models. The resulting predictions were then used to train SNAP (2013-02-16) through two rounds for final gene model predictions. Gene space completeness was assessed using BUSCO v3 and the eudicotyledons *odb10* pre-release dataset using standard parameters (Simão et al. 2015).

The predicted gene transcripts (mRNA) and predicted translated protein sequence were then annotated using Basic Local Alignment Search Tool Nucleotide (BLASTN) and Protein (BLASTP) 2.2.18+ for similarity to known transcripts and proteins, respectively. Alignments were made to the entire NCBI nucleotide and protein databases. For all BLAST homology searches, the e-value was set at 1e-25 and only the best match was considered. The predicted proteins were further annotated using InterProScan 5.28-67.0 for protein domain predictions (Mi et al. 2005; Camacho et al. 2009; Jones et al. 2014). InterProScan was run using standard settings. The complete assembly was analyzed using RepeatMasker 4.0.6 to search for small interspersed repeats, DNA transposon elements, and other known repetitive elements using the “Viridiplantae” repeat database and standard search parameters (Tarailo-Graovac and Chen 2009).

### Genomic Resequencing of Glyphosate Resistant Kochia and Differential Gene Expression

Genomic resequencing reads from the glyphosate resistant plant were aligned to the KoSco-1.0 genome assembly using the BWA-backtrack alignment program with default parameters (Li and Durbin 2009). The boundaries of the *EPSPS* copy number variant were manually detected where coverage dramatically increased up-and down-stream of the *EPSPS* gene.

RNA-Seq reads from susceptible and resistant plants were aligned to the gene models from the genome assembly using the mem algorithm from the BWA alignment program version 0.7.15 under standard parameters. Read counts for each gene were extracted from this alignment using the software featureCounts in the Subread 1.6.0 package and the gene annotation generated by WQ-Maker (Liao et al. 2013). Expression level and differential expression between the glyphosate susceptible and glyphosate resistant plants for all genes were calculated with the EdgeR package using the quasi-likelihood approach in the generalized linear model (glm) framework as described in the user manual (Robinson et al. 2010).

### Assembling the EPSPS Locus from a Glyphosate Resistant Plant

A library of bacteria artificial chromosomes (BACs) was generated from a single glyphosate resistant kochia plant selected from the glyphosate resistant population following the protocol described in Luo and Wing (2003) with modifications as described in Molin et al. (2017). High molecular weight (HMW) DNA was extracted from young leaf tissue from a single glyphosate resistant plant using a modified CTAB DNA extraction protocol. This HMW DNA was ligated to a linearized vector and transformed into *E. coli* using electroporation. Recombinant colonies were then grown on LB plates. Radiolabeled probes were designed for the *EPSPS* gene itself, a sequence upstream, and a sequence downstream of the *EPSPS* CNV. Predicted locations for the probes were determined by looking at the alignment of shotgun Illumina data from the glyphosate resistant line against the contig containing *EPSPS* in the genome assembly. Several colonies containing the appropriate sequences were identified for each probe. These identified BACs were end sequenced to determine their approximate location and run on pulse-field gel electrophoresis to determine their approximate size. Colonies containing positive BACs of the correct position and size were isolated and cultured. HMW DNA was extracted from these colonies and prepared using a SMRTbell Template Prep Kit, 1.0 using the manufacturer-recommended protocols. Finally, the HMW DNA was sent for RSII PacBio sequencing on two SMRT cells performed at The University of Delaware, DNA Sequencing & Genotyping Center.

PacBio reads were assembled using the software Canu (Koren et al. 2017). The BAC vector sequence was then removed from the assembled contigs. Using the known size of the BACs, their end-sequences and the corresponding contig from the susceptible genome assembly, entire BAC sequences were reconstructed manually from the contigs produced by CANU. These “full-length” BACs were then aligned, and overlaps were used to generate the largest contiguous length possible. This BAC meta-assembly was aligned to the susceptible contig from the genome assembly containing the *EPSPS* gene using YASS. Additionally, the BAC insert sequences were run through the MAKER pipeline, informed with cDNA and protein annotations from the Chenopodiaceae and the gene models from the kochia genome (Cantarel et al. 2008) for gene annotation. This BAC assembly led to the discovery of two dominant repeat types (a full length 56.1 kb repeat and a smaller 32.9 kb repeat), the upstream and downstream boundaries of the CNV, as well as a large mobile genetic element that was interspersed in the repeat structure.

Using the Illumina genomic resequencing data from the resistant line, we calculated the copy number of four regions from the CNV by read depth as follows: 1) the region directly upstream of the CNV; 2) the region directly downstream of the CNV; 3) the mobile genetic element; and 4) the full length, 56.1 kb repeat. This 56.1 kb repeat was then subdivided into the region only present within the 56.1 kb repeat and the region that is shared between the 56.1 kb repeat and a smaller 32.9 kb repeat. Highly repetitive regions and those containing transposable elements were masked for the alignment of resequencing reads. Genomic resequencing reads from the glyphosate resistant plant were aligned to these units using the BWA-backtrack alignment program using standard parameters. The number of reads mapping to each unit was calculated and divided by the length of that region to get the average number of reads per unmasked DNA length. The upstream and downstream read depths were averaged and used to standardize the read depths of each of the four units. These standardized read depths correspond with the predicted copy number of each unit.

### Markers for Confirming the Structure of the EPSPS CNV

Primers were designed that were spaced at regular intervals (~5 kb-15 kb) along the susceptible contig that spanned the putative CNV area for genomic qPCR analysis (Table 2). Additionally, qPCR primers were designed that spanned the junctions of the two dominant repeat types, the upstream and downstream boundaries of the CNV, as well as for the mobile genetic element (Table 2). Primers were designed to closely mimic the primers already published for the *EPSPS* gene (Wiersma et al. 2015), including a melting temperature between 51 and 56 °C, a GC content between 40 and 50%, and a length of between 20 and 24 bp. Furthermore, the resulting amplicon had to be between 100 and 200 bp long. All genomic PCR was performed using the same protocol established for *EPSPS* copy number assay (Gaines et al. 2016).

For genomic PCR screening of kochia populations for these repeat features, both susceptible and resistant plants were grown in the greenhouse until they were ~10 cm tall and 100 mg of young expanding leaf tissue was taken from each plant. DNA was extracted from this tissue using the recommended protocol from the DNeasy Plant Mini Kit. The DNA quality and concentration were checked using a NanoDrop 1000 and diluted to 5 ng/μl. For qPCR two genes were used as single-copy controls: acetolactate synthase (*ALS*) and copalyl di-phosphate synthetase 1 (*CPS*). Each qPCR reaction consisted of 12.5 μL PerfeCTa SYBR^®^ green Super Mix (Quanta Biosciences), 1 μL of the forward and reverse primers at 10 μM, 10 ng gDNA (2 μL), and 9.5 μL of sterile water for a total volume of 25 μL.

A BioRad CFX Connect Real-Time System was used for qPCR. The temperature cycle for all reactions was as follows: an initial 3 min at 95 °C followed by 35 rounds of 95 °C for 30 sec and 53 °C for 30 secs with a fluorescence reading at 497 nm after each round. A melt curve was performed from 65–95 °C in 0.5 °C increments for each reaction to verify the production of a single PCR product. Additionally, all products from a susceptible line were run on a 1.5% agarose gel to verify a single product with low to no primer dimerization. Relative quantification was calculated using the comparative Ct method: 2^ΔCt^ (ΔC_t_ = (Ct^(ALS)^+Ct^(CPS)^)/2 − C_t_^EPSPS^) (Schmittgen and Livak 2008).

### Data Access

All raw sequence read files for the whole genome sequencing have been deposited in the Sequence Read Archive database at NCBI under BioProject ID PRJNA526487 (SRR8835960-SRR8835963). The genome assembly was submitted to the NCBI genomes database with the accession SNQN00000000.

## RESULTS

### Genome Assembly and Annotation

The KoSco-1.0 assembly consisted of 19,671 scaffolds, spanning 711 Mb. The longest scaffold was 770 kb and the N50 was 62 kb for this assembly. Approximately 9.43% of the base pairs were unknown “N” bases that serve only as scaffolding and distance information (Sup. Table 1). After annotation with Maker, 47,414 genes were predicted in KoSco-1.0 with an average transcript length of 943 bp (Sup. Table 2), compared to the 27,429 genes in *Beta vulgaris* (Dohm et al. 2014). KoSco-1.0 was analyzed using BUSCO for completeness, which found 1,490 out of 2,121 (70.3%) ultra-conserved genes from the eudicotyledons *odb10* dataset (Sup. Table 3). Approximately 62% of predicted kochia genes found one or more matches in the NCBI database(s) using a BLAST e-value < 1e-25 and almost 82% of predicted proteins were assigned one or more functional InterPro domain(s) (Sup. Table 2). RepeatMasker uncovered 6.25% of the genome assembly consisting of interspersed repeats with the largest proportion consisting of LTR elements of either the Ty1/Copia or Gypsy/DIRS1 variety. Simple repeats made up approximately 2.5% of the assembly (Sup. Table 4).

**Table 1.** List of genes near *EPSPS* that are in or flanking the *EPSPS* CNV event. Read depth is the log_2_ of the difference between the background read depth and the read depth of each gene from genomic Illumina sequencing of a glyphosate resistant line. Base-pair coordinates are given relative to their position in the contig from the susceptible genome assembly. DE is the log_2_ differential expression between four resistant and four susceptible individuals from RNA-Seq. P-value is the significance of DE and is adjusted for false discovery rate.

| Gene | Beginning | Ending | Length | Orientation | Description | Part of the CNV? | Read Depth | DE | P-value |
| --- | --- | --- | --- | --- | --- | --- | --- | --- | --- |
| KS_00451 | 27,406 | 28,674 | 1,268 | Reverse | GRAVITROPIC IN THE LIGHT 1-like | No | 0 | -0.43 | 0.00 |
| KS_00452 | 35,728 | 36,696 | 968 | Reverse | IRK-Interacting Protein | No | 0 | -2.62 | 0.05 |
| KS_00453 | 37,839 | 41,640 | 3,801 | Reverse | Nitroreductase family | No | 0 | 0.74 | 0.00 |
| KS_00454 | 43,124 | 47,121 | 3,997 | Forward | arginase 1, mitochondrial | 56.1 kb | 2.86 | 2.23 | 0.00 |
| KS_00455 | 47,240 | 52,651 | 5,411 | Reverse | protein NRT1/ PTR FAMILY 7.2-like | 56.1 kb | 2.86 | 0.72 | 0.58 |
| KS_00456 | 63,014 | 72,467 | 9,453 | Forward | tRNA N6-adenosine threonylcarbamoyltransferase | 56.1 kb & 32.7 kb | 3.49 | 3.03 | 0.00 |
| KS_00457 | 72,617 | 73,531 | 914 | Reverse | golgin subfamily A member 6-like | 56.1 kb & 32.7 kb | 3.49 | -3.18 | 0.00 |
| KS_00458 | 76,342 | 81,181 | 4,839 | Forward | DNA repair protein RAD51 | 56.1 kb & 32.7 kb | 3.46 | 1.33 | 0.00 |
| KS_00459 | 82,421 | 84,836 | 2,415 | Forward | transketolase, chloroplastic-like | 56.1 kb & 32.7 kb | 3.29 | 3.83 | 0.00 |
| KS_00460 | 91,663 | 97,214 | 5,551 | Forward | 3-phosphoshikimate 1-carboxyvinyltransferase 2 (EPSPS) | 56.1 kb & 32.7 kb | 3.12 | 4.01 | 0.00 |
| KS_00461 | 106,901 | 109,241 | 2,340 | Forward | NAD dependent epimerase | No | 0 | 2.52 | 0.00 |
| KS_00462 | 106,975 | 110,332 | 3,357 | Reverse | uncharacterized protein | No | 0 | 2.54 | 0.06 |
| KS_00463 | 113,504 | 114,006 | 502 | Reverse | DUF861 | No | 0 | 0.05 | 0.85 |

**Table 2.**
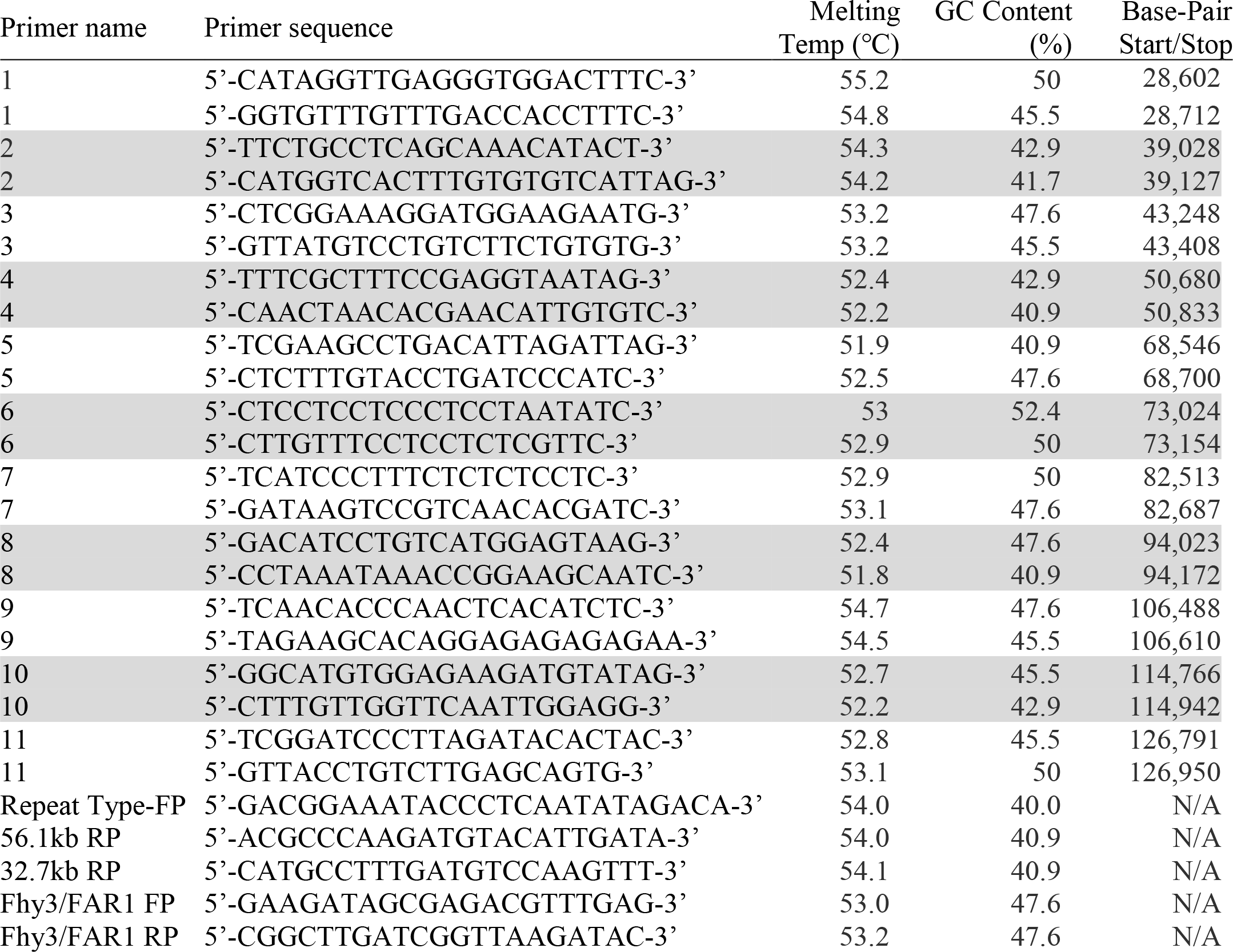
Primers for qPCR markers for determining copy number at multiple locations near the *EPSPS* gene and qPCR markers for determining copy number of 56.1 kb repeats, 32.7 kb repeats, and the MGE. Base-pair coordinates of PCR amplicons are given relative to their position in the contig from the susceptible genome assembly

**Table 3.**
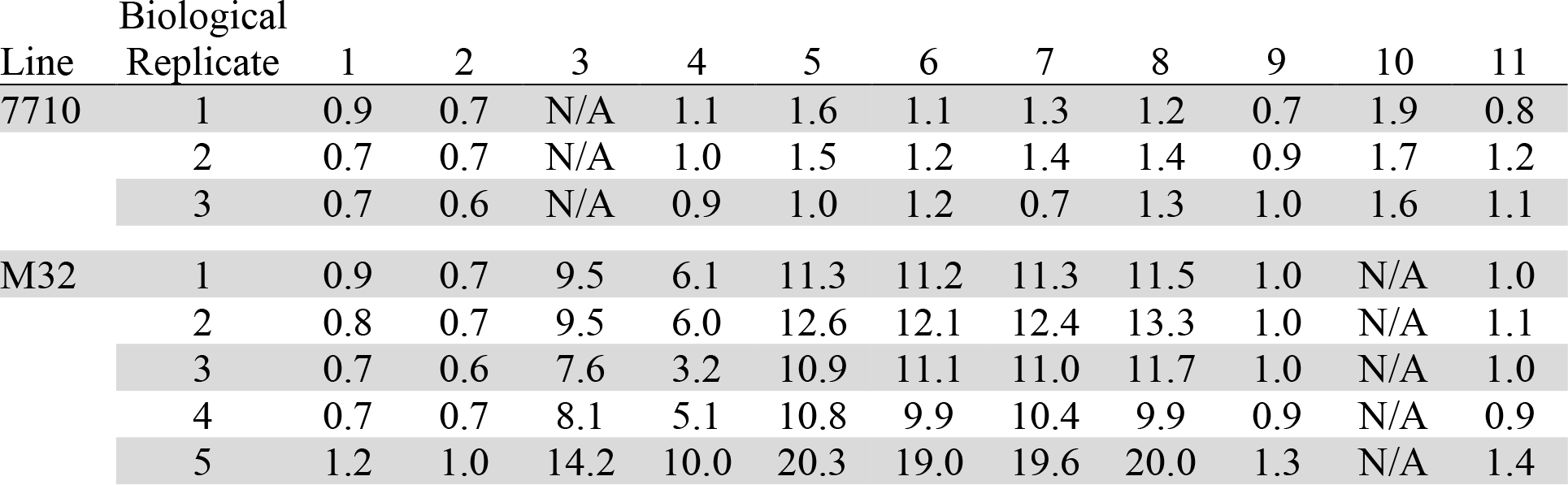
Copy number data from all qPCR markers on three glyphosate-susceptible (7710) and five glyphosate-resistant (M32) individuals. Copy number is calculated as ΔC_t_= (C_t_^(ALS)^+C_t_^(CPS)^)/2 − C_t_^Marker^. “N/A” stands for “No Amplification”.

**Table 4.** Copy number data for the number of 56.1 kb repeats, 32.7 kb repeats, and the MGE on three glyphosate-susceptible (7710) and five glyphosate-resistant (M32) individuals. Copy number is calculated as ΔC_t_= (C_t_^(ALS)^+C_t_^(CPS)^)/2 − C_t_^Marker^. “N/A” stands for “No Amplification”.

| Line | Replicate | 56.1kb | 32.7kb | MGE |
| --- | --- | --- | --- | --- |
| 7710 | 1 | N/A | N/A | 3.9 |
|  | 2 | N/A | N/A | 5.5 |
|  | 3 | N/A | N/A | 4.7 |
| M32 | 1 | 5.4 | 1.8 | 16.2 |
|  | 2 | 5.1 | 1.9 | 17.4 |
|  | 3 | 5.1 | 1.7 | 18.2 |
|  | 4 | 5.3 | 1.7 | 14.1 |
|  | 5 | 6.9 | 2.1 | 17.7 |

### The EPSPS Locus and Differential Gene Expression

The contig containing the *EPSPS* locus from the susceptible genome assembly was 399,779 bp long. The *EPSPS* gene model was 5,551 bp long (UTRs, exons, and introns included) and located between base pairs 91,663-97,214 of the contig. When this contig was aligned to *Beta vulgaris* near perfect synteny was observed; however, when compared to the sequence responsible for duplicating *EPSPS* from *Amaranthus palmeri*, little similarity existed outside of the *EPSPS* gene itself (Figure 1).

**Figure 1.**
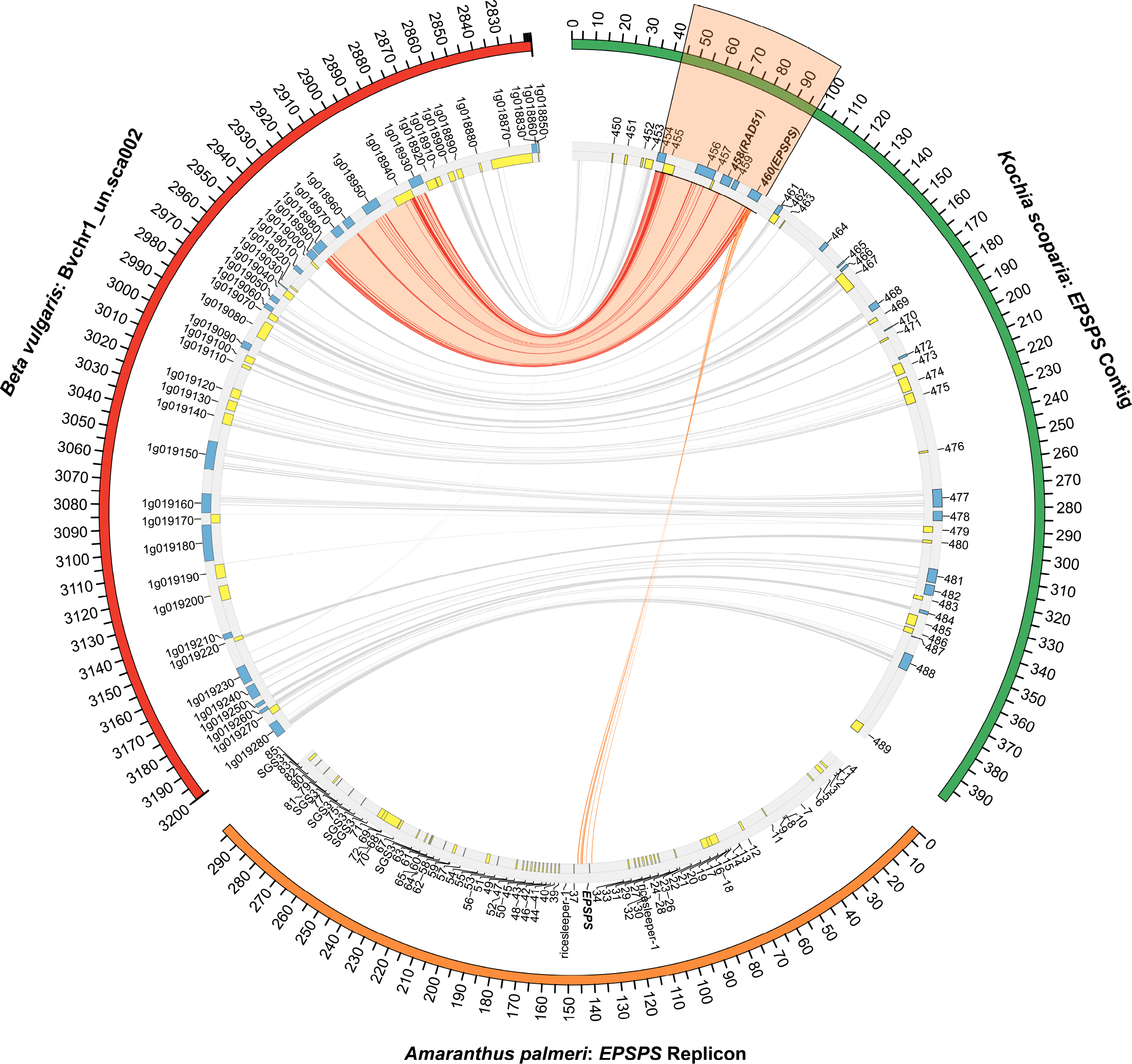
A comparison of the *EPSPS* contig from kochia (Green), an genomic scaffold from chromosome 1 of the *Beta vulgaris* genome (Red) (Genbank ID: KQ090199.1) (Dohm et al. 2014), and the *EPSPS* replicon from *Amaranthus palmeri* (Orange) (Molin et al. 2017). Blue and yellow blocks indicate genes in the forward and reverse orientation, respectively. The *EPSPS* gene is highlighted in orange. Red, connecting lines, indicate areas of high similarity between *Beta vulgaris* and kochia. Orange, connecting lines indicate areas of high similarity between *Amaranthus palmeri* and kochia. Number of base pairs in the alignment are listed on the outside track. The links between *Beta vulgaris* and kochia that fall within the *EPSPS* duplicated region are highlighted in orange.

When shotgun Illumina genomic reads from the glyphosate resistant line were aligned to the contig, the read depth of *EPSPS* and its surrounding area was much greater (> 7.26-fold) than the background read depth. Using this alignment, it was possible to predict the exact boundaries of the *EPSPS* CNV starting at base pair 41,684 and continuing to base pair 101,128. This region contains seven coding genes of various functions including *EPSPS* itself (Table 1). When differential expression of all genes in the genome was calculated using RNA-Seq data, five of the genes in this region showed over expression in the glyphosate resistant line, one gene showed under-expression in the glyphosate resistant line, and one showed no significant difference (FDR adjusted p-value < 0.05) (Table 1). Since gene expression is dynamic, depending on both environmental conditions and developmental stage, the genes not showing DE may be overexpressed in glyphosate resistant plants under different experimental conditions. When the *EPSPS* contig was aligned to itself, there was no evidence for sequence complexity (simple sequence repeats, inverted repeats, self-homology, etc.) at the predicted boundaries of the CNV (Sup. Figure 2).

**Figure 2.**
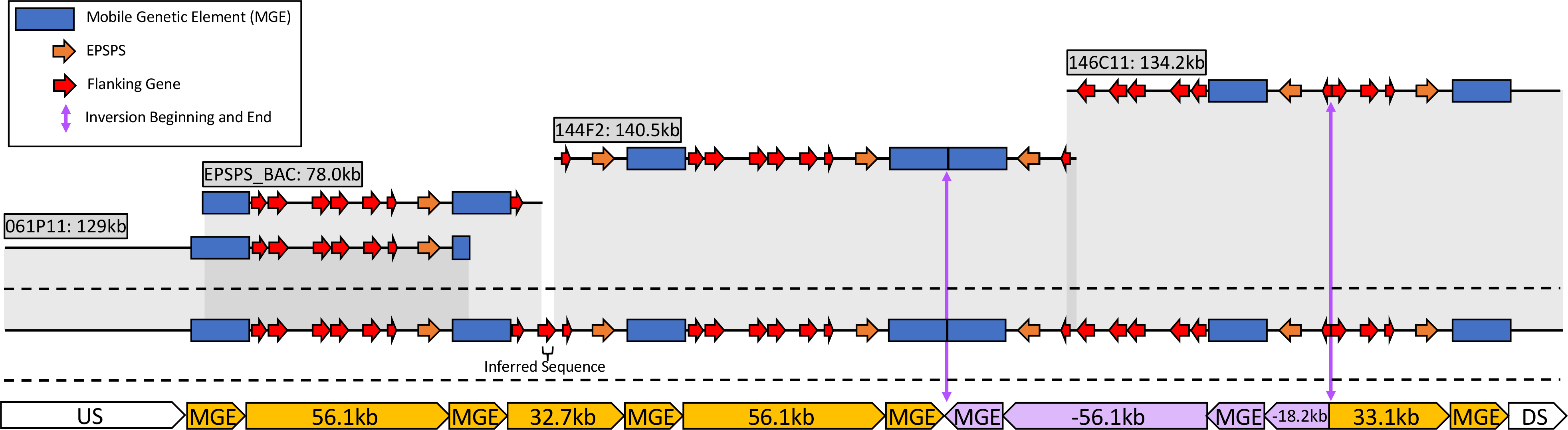
A diagram of the four assembled BACs and how they overlap to generate five different repeat types of the *EPSPS* CNV locus from glyphosate resistant kochia. The mobile genetic element (MGE) is illustrated as a blue rectangle, the *EPSPS* gene is a green arrow, the co-duplicated genes are orange arrows, and the beginning and end of the inverted repeat are vertical arrow lines.

### The EPSPS Locus from a Glyphosate Resistant Plant

Using PacBio data of four BACs from a glyphosate resistant plant, we assembled four contigs that were 129.0 kb for the BAC detected with the upstream probe, 134.2 kb for the BAC detected with the downstream probe, and 140.5 kb and 78.0 kb for two BACs detected with the *EPSPS* probe. These assemblies encompassed at least six repeats of the *EPSPS* gene and a significant portion of the upstream and downstream sequence. The largest and most complete repeat was 56.1 kb long and contained the entire region predicted from the alignment of resistant Illumina data against the susceptible *EPSPS* contig, including all seven of the predicted genes in this region. The second type was 32.7 kb and contained only four of the seven co-duplicated genes from the 56.1 kb repeat, including *EPSPS* and the three genes immediately upstream of it. The third repeat was a full-length inversion of the 56.1 kb repeat. The fourth type of repeat was an 18.2 kb inverted repeat that contained only *EPSPS* and a fraction of one upstream gene. The fifth and final repeat structure was identified as a forward repeat of 33.1 kb, containing *EPSPS* and the three genes immediately upstream of it (Figure 2). All repeats end at the same downstream base pair, directly after *EPSPS*; however, the beginning upstream base pair of each repeat type is variable (Figures 2 and 3).

**Figure 3.**
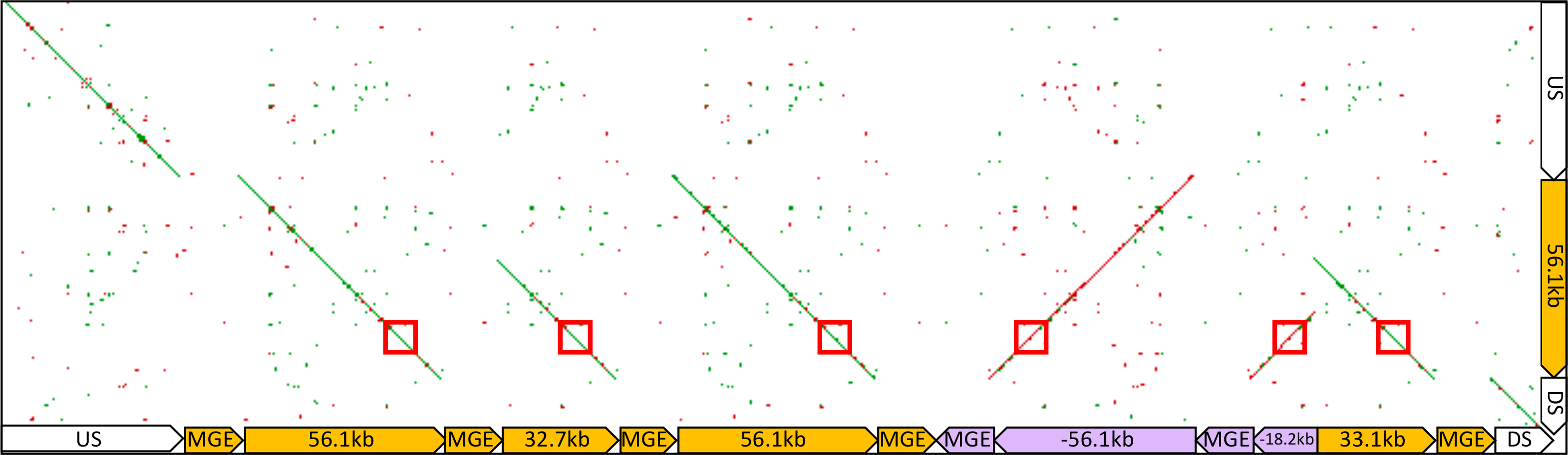
A dot-plot alignment of the assembled resistant *EPSPS* locus to the contig containing *EPSPS* from the susceptible genome assembly. The location of *EPSPS* is indicated by a red box. Large gaps in alignment are the insertion sites of the MGE.

Enough overlap existed among the BAC contigs to composite all BAC assemblies together to make a representative sequence (meta-assembly) that contained two full-length 56.1 kb repeats and one of each of the other repeat types. Additionally, the flanking single-copy upstream and downstream sequences were included. When this BAC meta-assembly from glyphosate resistant kochia was aligned to the susceptible contig from the genome assembly, we observed perfect agreement between the resistant and susceptible loci; however, a large disparity was evident at each repeat junction and on either end of the resistant repeat structure (Figures 3). A 16,037 bp sequence was inserted just downstream and upstream of all repeats in the glyphosate resistant BAC assemblies. This insert shows no homology with any part of the susceptible contig; furthermore, when this insertion was aligned against the entire susceptible genome, this region was not found in its entirety.

Maker was run on this insertion to predict gene models and identified four regions with putative coding genes. The first predicted gene belonged to the family of genes known as FHY3/FAR1 (IPR031052) and contained the domains: “AR1 DNA binding” and “zinc finger, SWIM-type” (IPR004330F, IPR007527 respectively). The second gene’s function was less clear but was identified to be part of the Ubiquitin-like domain superfamily (IPR029071). The third gene’s function was also unclear and was generally identified as belonging to the Endonuclease/exonuclease/phosphatase superfamily (IPR036691). The fourth and final gene had no identifiable InterPro domains, and had BLAST hits to uncharacterized proteins in NCBI. We refer to this insertion as the mobile genetic element (MGE) in all figures and discussion as it seems to have inserted only in resistant lines from an unknown *trans* location in the genome.

### Markers for Confirming the Structure of the EPSPS CNV

Quantitative PCR markers were developed dispersed across the entire CNV, including markers on both sides in regions that show no evidence of CNV (Table 3). These markers performed, for the most part, as predicted based on the resequencing of the glyphosate resistant plants and the BAC sequencing. All markers upstream and downstream of the CNV are approximately single copy. Markers 3 and 4, predicted to be only in the longer, 56.1 kb repeat, both show increased copy number in resistant individuals. Markers 5, 6, 7, and 8, are in both 56.1 kb and 32.7 kb repeats. These four markers were tightly associated, co-varied for each individual, and showed higher copy number than markers 3 and 4 (Table 3).

Additional qPCR markers were developed that only amplified when the MGE was flanked by either the two dominant repeat types of 56.1 kb or 32.7 kb. Using these markers, we quantified the number of 56.1 kb or 32.7 kb repeats in several individuals. In our line, 32.7 kb repeats were less frequent then 56.1 kb repeats. The tested individuals each had approximately two 32.7 kb repeats and between five and seven 56.1 kb repeats (Table 4). These markers did not amplify in any susceptible plants, which supports the discovery that the MGE is not present at the beginning of the susceptible *EPSPS* locus.

Additionally, we developed a marker internal to the MGE. All susceptible individuals had approximately 4-5 copies of this marker; however, none of these regions were present in the KoSco-1.0 genome assembly. In resistant individuals, we detected 14-18 copies of the MGE. If we account for the 4-5 copies that are in the susceptible individuals and if we consider that a MGE exists at both the upstream and downstream boundary, then we would predict 9-13 copies, which almost perfectly correlates with the copy number observed for qPCR markers 5, 6, 7, and 8. This would indicate that one copy of the MGE is associated with each repeat (Table 4).

Illumina shotgun genome resequencing data from a resistant kochia plant aligned to four distinct units from the BAC assembly was used to calculate the copy number of each unit of the repeat structure and to confirm our qPCR results. After standardizing the read depth of each unit by the background read depth, we calculated 7.4 copies of the 56.1 kb repeat, 10.9 copies of the 32.7 kb repeat type, and 14.3 copies of the mobile genetic element (Figure 4A, 4B). It should be noted that the unit of the 32.7 kb repeat type includes reads from all repeats due to the sequence of this region being shared in all repeat types. With this information in conjunction with previously published cytogenetic work (Jugulam et al. 2014; Jugulam and Gill 2018), we propose a model for the structure of the *EPSPS* CNV from resistant kochia individuals (Figure 5).

**Figure 4.**
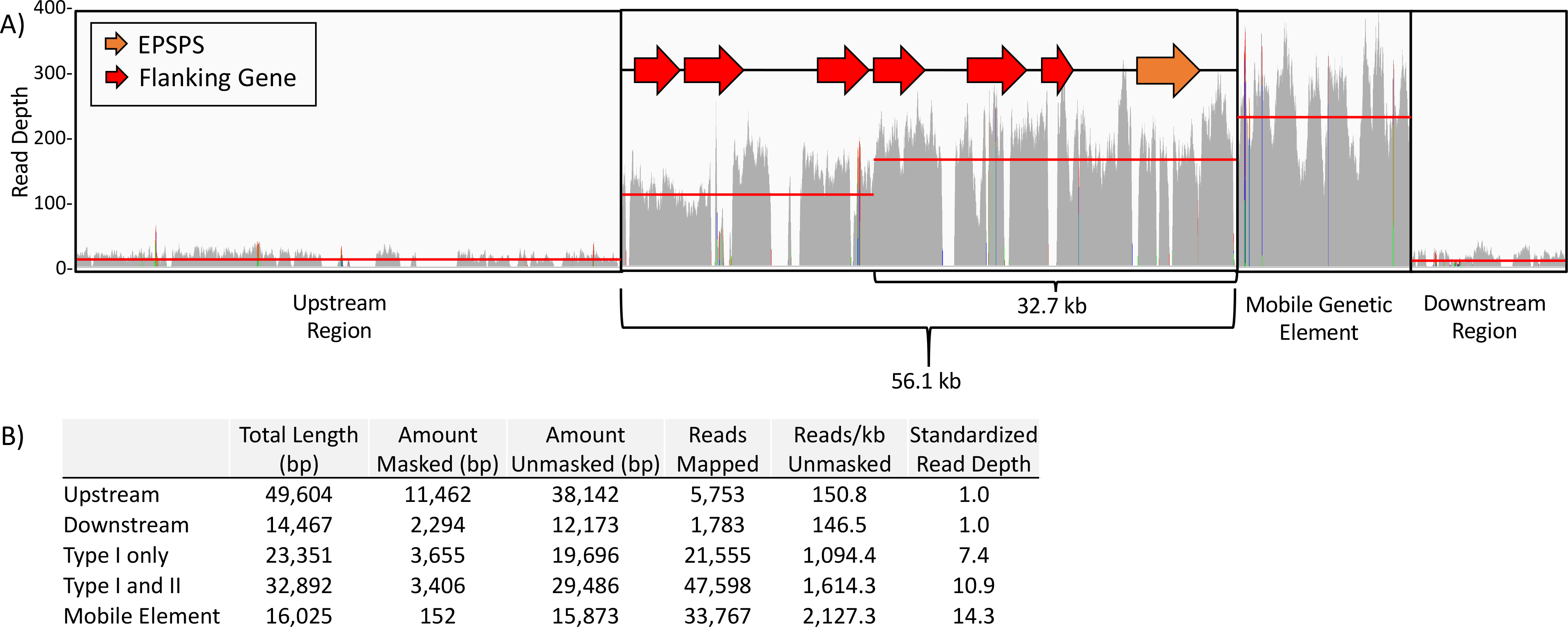
A) Illumina shotgun genome resequencing data from a resistant kochia plant aligned to four distinct units from the BAC assembly: 1) The region directly upstream of the *EPSPS* tandem duplication, 2) the tandemly duplicated region of the genome containing *EPSPS*, 3) the MGE, and 4) the region directly downstream of the *EPSPS* tandem duplication. Red lines indicate the average read depth for that unit. Two averages are indicated for the tandemly duplicated region of the genome containing *EPSPS* due to two major repeat sites existing in the *EPSPS* CNV structure: the 56.1 kb and 32.7 kb repeat types. B) A table outlining the calculation for copy number estimates for the four units. The total length of the region, the amount of repetitive DNA that was masked, the amount of DNA remaining unmasking, the number of reads mapped to the unmasked regions, the average reads per kilobase of unmasked DNA, and the read depth divided by the reads/kb unmasked of the non-duplicated region.

**Figure 5.**
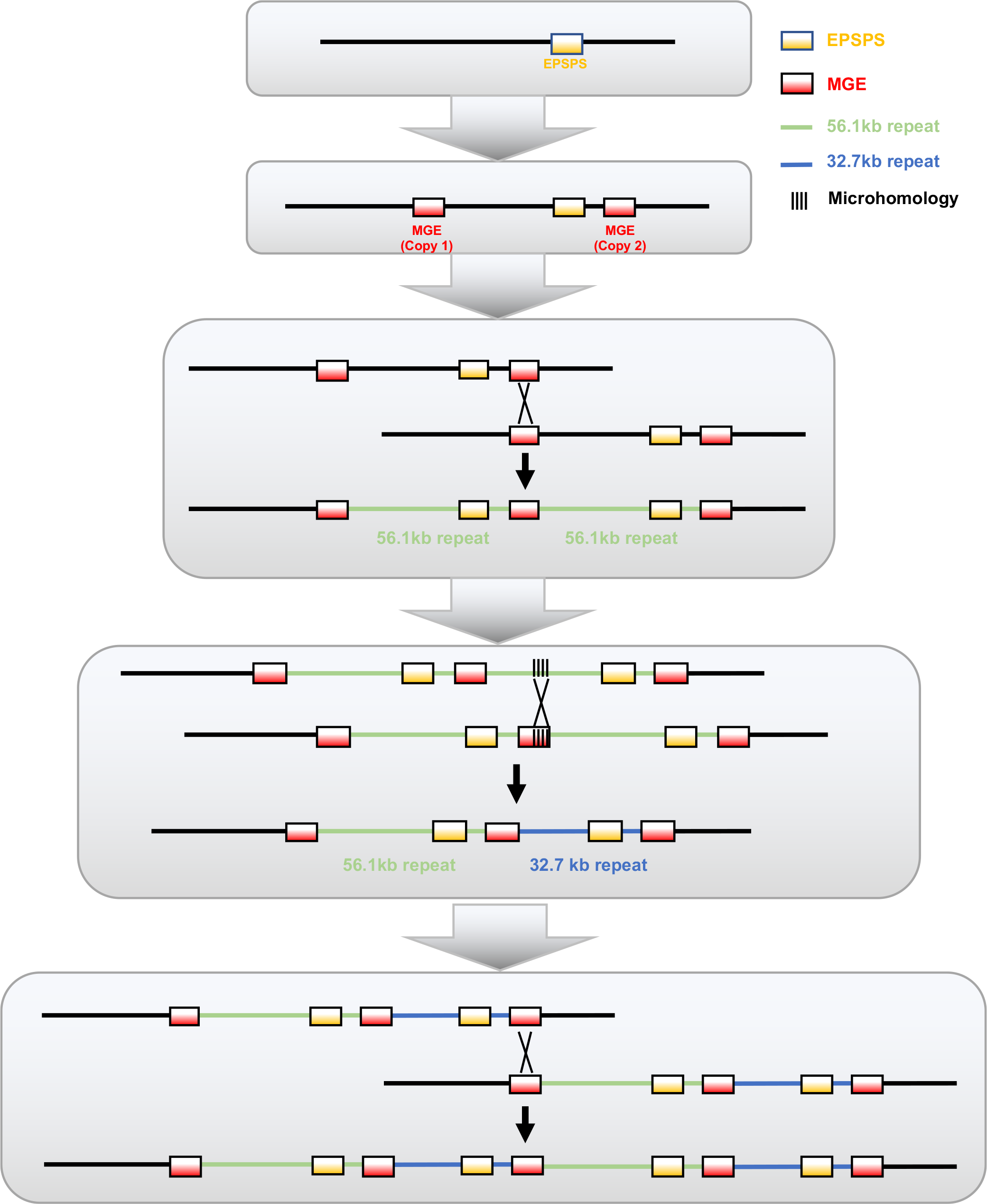
A model for the generation and continued increase of *EPSPS* copy number. The initial event that led to *EPSPS* gene duplication was the insertion of two mobile elements both upstream and downstream of the *EPSPS* gene (MGE). After unequal crossing over, gametes were produced with >1 *EPSPS* gene copy. Subsequently, a double stranded break occurred at the MGE boundary that was incorrectly repaired using a microhomology-mediated mechanism within the middle of the repeat region, generating a shorter copy of this repeat region (32.7kb repeat).

## DISCUSSION

### Structure and Genetic Content of the EPSPS Tandem Duplication Region

The *EPSPS* contig from susceptible kochia has near perfect synteny with *Beta vulgaris* along its entire length but little homology with the *EPSPS* region from *Amaranthus palmeri* (Figure 1). Thus, multiple species within the Caryophyllales have independently evolved glyphosate resistance via *EPSPS* gene duplication but have done so through very different genomic mechanisms: tandem duplication vs. proliferation of an extrachromosomal element (Jugulam et al. 2014; Molin et al. 2017; Koo et al. 2018; Patterson et al. 2018).

We discovered the genomic elements that constitute the two most dominant repeats in the tandem duplication. Additionally, we discovered a MGE in between each repeat. Taking everything into account, there is most often either 72.6 kb or 49.2 kb between *EPSPS* genes in the CNV locus. These estimates are similar to but slightly larger than the previously fiber-FISH estimated sizes of 66 kb and 45 kb respectively in another resistant kochia line (Jugulam et al. 2014). What accounts for the differences between our assemblies and the previously reported fiber-FISH studies remains unclear, as Fiber-FISH can have a resolution of ~1 kb (Ersfeld 1994). It may be that different populations of kochia have different repeat sizes. Further testing and validation on the type and size of the *EPSPS* repeats in various, divergent populations is needed to confirm this. We did detect an inverted repeat near the downstream end of the CNV as shown by Jugulam et al. (2014).

RNA-Seq expression data shows that four of the six genes within the conserved region of the tandem-repeat are over-expressed at a rate commensurate with genomic resequencing read depth: *RAD51*, *transketolase*, *tRNA N6-adenosine threonylcarbamoyltransferase*, and *EPSPS* (FDR adjusted p-value <0.05). The expression of two other genes (*golgin subfamily A member 6-like protein 6* and *NRT1/ PTR Family 7.2-like*) is reduced in the resistant line and may be due to gene silencing, similar to what happens when multiple copies of transgenes are inserted in the same plant (Finnegan and McElroy 1994; Tang et al. 2006) (Table 1). The obvious benefit of *EPSPS* over-expression is glyphosate resistance, but the phenotypic effects due to increased expression of other genes in this CNV remain unclear.

The expression of the *RAD51* homolog is especially interesting due to its importance in regulating crossing over. Mis-expression, up or down, of *RAD51* has been shown to cause cancer in animal tissues as *RAD51* is involved in regulating homologous recombination of DNA during double stranded break repair (Maacke et al. 2000) (Table 1). Additionally, RAD51, along with the recombinase DMC1, facilitate recombination of homologous chromosomes during meiosis in plants and animals (Crickard et al. 2018). In humans, *RAD51* expression is modulated by miRNAs and mis-regulation of these miRNAs are often associated with various forms of cancer (Choi et al. 2014; Gasparini et al. 2014; Cortez et al. 2015; Liu et al. 2015a; Liu et al. 2015b). Therefore, we would predict that over-expression of *RAD51* in the resistant line would have a large impact phenotypic consequence and could change the recombination rates and double strand break repair.

We used qPCR genomic copy number primers to validate much of our BAC assembly. The results from a pair of primers that detected the presence and number of the MGE were surprising. In the susceptible plant, approximately 4-6 MGE copies were observed despite not appearing in the susceptible genome assembly; therefore, this MGE is present in the susceptible plant but it was not assembled in the whole genome assembly. It may be that these background copies lie in repetitive or difficult to assemble regions. In the resistant plants, the number of MGE copies was always approximately equal to the *EPSPS* copy number plus 4-6 copies, indicating that the original copies found elsewhere in the genome are still present and the insert is being co-duplicated with every repeat of the *EPSPS* CNV. The fact that the MGE also seems to be in the susceptible lineage implies that the insertion in the *EPSPS* region originated by transposition within the genome.

### The Role of a Mobile Genetic Element in EPSPS Gene Duplication

When the *EPSPS* contig from the susceptible genome assembly is aligned to itself, no complexities, such as SSRs or large homodimers of nucleotides, exist at the beginnings of any of the repeat types (Sup. Figure 1). This would indicate that the sequence in the susceptible locus alone is insufficient for explaining why this region has become a site for copy number variation, which is inconsistent with earlier predictions that homology exists at the upstream and downstream boundaries where an initial misalignment occurred (Jugulam et al. (2014). Mobile genetic elements, such as transposons, have been proposed to cause tandem repeats of sequences near their insertion point (Tsubota et al. 1989; Reams and Roth 2015).

We propose that the insertion of a MGE near the *EPSPS* locus in the resistant kochia line facilitated the subsequent history of tandem duplication in this region. The MGE contains a member of the Fhy3/FAR1 gene family. Genes in this family are thought to be derived from MULE transposons and have been “domesticated” to have a role in the regulation of genes involved in circadian rhythm and light sensing in a wide phylogentic distribution of angiosperms (Wang and Deng 2002; Hudson et al. 2003; Cowan et al. 2005; Tang et al. 2012). We hypothesize the insertion of the MGE near the *EPSPS* locus in resistant kochia line is evidence that Fhy3/FAR1 elements may still be mobile and that they are not fully “domesticated.” Because the insert appears to be both at the upstream and downstream borders of the CNV, we hypothesize that insertions of this MGE happened in two locations, flanking the *EPSPS* region. These two insertions then could have led to misalignment during meiosis as both MGEs are identical. A subsequent crossing-over event somewhere along the length of the misaligned MGE copies would have generated two alleles – one with two of the more common 56.1 kb repeats, and the other with no *EPSPS* gene, the latter of which would be lethal in the homozygous state. Such unequal crossing over could then facilitate further expansions of this region.

Interestingly, the MGE boundary shares 7-bp of sequence identity with the precise beginning of the shorter, less common 32.7 kb repeat. We propose that a recombination event took place between the MGE downstream boundary and the start site of the smaller 32.7 kb repeat, perhaps mediated by double-stranded break repair at the end of the MGE (Figure 5) (Ottaviani et al. 2014; Sfeir and Symington 2015). Short microhomology-mediated illegitimate recombination has been well studied in bacteria (Petes and Hill 1988; Nash 1996; Romero and Palacios 1997; de Vries and Wackernagel 2002; Reams and Neidle 2004). The presence of the MGE end at the breakpoint of the large inversion in the tandem array (Figure 2) further implicates double-stranded breaks at the MGE boundaries with the genome instability in this region. Homologous recombination and double strand break repair depend heavily on the enzyme RecA in bacteria and its homologue RAD51 in eukaryotes. These enzymes bind single-stranded DNA and promote strand invasion and therefore the exchange between homologous DNA molecules (Baumann and West 1998; Lin et al. 2006; Hastings et al. 2009). In kochia, it remains unclear if the presence of *RAD51* in the duplicated region is coincidental or has affected the evolution of this tandem duplication event.

### EPSPS Duplication in Weeds

Cytological evidence in kochia has previously shown that *EPSPS* gene duplication in kochia was due to tandem duplication and not by *trans*-duplication (Jugulam et al., 2014). However, this work was limited to cytology and was unable to pinpoint the sequence differences between resistant and susceptible plants. Additionally, the genetic content of the region outside of the *EPSPS* gene was unknown. Our work has resolved these uncertainties, providing a clearer understanding of the structure of the *EPSPS* gene duplication event and enabling investigation of the exact phenotypic and evolutionary consequences of this event.

Eight plant species have been confirmed to have evolved resistance to glyphosate via increased *EPSPS* gene copy number (reviewed in Patterson et al. 2018). Of these, only the genetic mechanisms of gene duplication in *Amaranthus palmeri* have been investigated and explained. In the case of *Amaranthus palmeri*, duplicated *EPSPS* genes are carried on a large extra-chromosomal circular DNA that is inherited by tethering to the chromatin (Molin et al., 2017, Koo et al., 2018). Kochia and *Amaranthus palmeri* are both members of the Caryophyllales; however, each species has independently evolved glyphosate resistance by *EPSPS* gene duplication from completely different genetic mechanisms.

## CONCLUSION

Widespread and repeated use of the herbicide glyphosate represents an intense abiotic selective pressure across large areas. Several weed species have evolved resistance to this pressure by means of increased copies of the target-site gene *EPSPS*. We identified a MGE at the duplicated *EPSPS* locus and hypothesize that the insertion of one or more of these MGEs initiated a tandem duplication event. Once the initial gene duplication occurred, the locus had unequal recombination producing gametes with increased and decreased copy numbers. This interplay between transposable elements and target site copy number variation provides valuable insight into how genomic plasticity may contribute to rapid evolution of abiotic stress tolerance. Continuing to investigate the roles transposable elements and gene duplication play in shaping plant resilience is essential for understanding evolution and how plant genomes are changing in response to human activities.

## ACKNOWLEDGEMENTS

This work was partially supported by the Colorado Wheat Administrative Committee, Dow AgroSciences, and by the USDA National Institute of Food and Agriculture, Hatch project COL00783, accession number 1016207, to the Colorado State University Agricultural Experiment Station.

