## Supporting Information for "The Draft Genome of *Kochia scoparia* and the Mechanism of Glyphosate Resistance via Transposon-Mediated *EPSPS* Tandem Gene Duplication"

Methods

DNA Extraction

For shotgun genome Illumina sequencing of the two lines, DNA was extracted from samples using a modified CTAB extraction protocol described in Doyle (1991). First, 500 µl of extraction buffer (100 mM tris, 1.4 M NaCl, 20 mM EDTA, pH 8.0, 2% CTAB, 0.3% mercaptoethanol) with 5 mg polyvinylpyrrolidone (PVP) was mixed with the tissue aliquots. The suspension was homogenized and incubated at 60 °C for 15 min. Next, 500 µl of chloroform:isoamyl alcohol (24:1) was added and the tubes were gently agitated on an orbital mixer for 15 min. The tubes were then centrifuged at 8000 rcf for 15 min and the top, aqueous phase was moved to a new tube. One µl of RNase A was added and incubated at 37 °C for 1 hour. The chloroform:isoamyl alcohol separation was performed again and the aqueous phase retained again. DNA was then precipitated by adding 1/10 volume 5M sodium acetate, pH 8 and three volumes of 100% ethanol. The samples were then centrifuged at 10,000 rcf for 10 min. The supernatant was poured off and the resulting pellet was rinsed with 70% ethanol and then allowed to dry. The pellet was re-suspended in 100 µl of water, checked for concentration and purity on a Nanodrop T1000.

For large-fragment, genomic PacBio sequencing of the glyphosate-susceptible line, the CTAB protocol was further modified to obtain more DNA of sufficiently large size (>10kb). Approximately 1 g of finely chopped kochia young leaf tissue was added to 50 ml conical tubes. To this tissue 15 ml of CTAB extraction buffer and 60 µg of PVP were added, mixed, and allowed to incubate for 30 min at 50 °C. The tubes were then centrifuged at 3600 rcf for 10 min. The liquid phase was separated into a new tube and 15 mL of chloroform:isoamyl alcohol (24:1) was added and mixed by inversion. They were then centrifuged at 3600 rcf for 10 min more and the upper phase transferred to a new tube. To this 4 µl of RNase A was added and incubated for 30 min at 37 ℃. The chloroform:isoamyl alcohol separation was repeated and the final aqueous phase collected. The DNA was precipitated by adding 3 volumes of EtOH and 0.5 volume of NaCl 5M. The tubes were then incubated at -20 ℃ for 30 min, centrifuged at 3600 rcf for 10 min, and the pellet washed with 70% ethanol. The final pellets were dried and re-suspended in 2 ml of Tris-EDTA buffer. The DNA was further purified using the Genomic DNA Clean & Concentrator™-10 kit by Zymo, following the recommended protocol. The DNA was checked for quality using a NanoDrop 2000c and quantified using Qubit.
